# Neural correlates of effort-based valuation with prospective choices

**DOI:** 10.1101/357327

**Authors:** Nadav Aridan, Nicholas J. Malecek, Russell A. Poldrack, Tom Schonberg

## Abstract

How is effort integrated in value-based decision-making? Animal models and human neuroimaging studies, primarily linked the anterior cingulate cortex (ACC) and ventral striatum (VS) to the integration of effort in valuation. Other studies demonstrated the role of these regions in invigoration to effort demands, thus it is hard to separate the neural activity linked to anticipation and subjective valuation from actual performance. Here, we studied the neural basis of effort valuation separated from performance. We scanned forty participants with fMRI and they were asked to accept or reject monetary gambles that could be resolved with future performance of a familiar grip force effort challenge or a fixed risk prospect. Participants’ willingness to accept prospective gambles reflected discounting of values by physical effort and risk. Choice-locked neural activation in contralateral primary sensory cortex and ventromedial prefrontal cortex (vmPFC) tracked the magnitude of prospective effort the participants faced, independent of choice time and monetary stakes. Estimates of subjective value discounted by effort were found to be tracked by the activation of a network of regions common to valuation under risk and delay, including vmPFC, VS and sensorimotor cortex. Together, our findings show separate neural mechanisms underlying prospective effort and actual effort performance.

## Introduction

Effortful behavior, whether among laboratory participants or foraging animals, generally demonstrates that effort imposes a cost on valuation, revealed by discounting of rewards and preferences for less effortful alternatives (Hull 1943; Charnov 1976; Rudebeck et al. 2006). Theoretical models of valuation suggest that the integration of expected cost and benefit representations guides choice behaviors (Rangel et al. 2008; Padoa-Schioppa 2011). Converging evidence from animal models and human neuroimaging studies link integrated value to neural activity within a network of regions that includes the ventral striatum (VS), posterior cingulate cortex, anterior insula and ventromedial prefrontal cortex (vmPFC; Bartra, McGuire, & Kable, 2013; Clithero & Rangel, 2013; Rushworth & Behrens, 2008) and demonstrate modulation of its activity by common costs such as risk (Mohr et al. 2010) and delay (Carter et al. 2010). However, accounts of effort-based valuation have instead emphasized the specific role of the anterior cingulate cortex (ACC) together with VS. For example, animals with lesions to ACC (Walton et al. 2003; Rudebeck et al. 2006), or depletion of dopamine within VS (Salamone et al. 2007) fail to allocate effort properly to maximize available rewards. Patterns of phasic dopamine release within VS (Day et al. 2010) and the activity of single neurons within ACC reflect the association of effort cost to expected reward in isolation (Pasquereau and Turner 2013) and competitive contexts (Hillman and Bilkey 2012). Human neuroimaging studies extended the relationship of VS to effort-discounted reward (Croxson et al. 2009; Kurniawan et al. 2010) and the ACC to effort-based cost-benefit analyses (Prévost et al. 2010; Kurniawan et al. 2013). Related studies also implicated primary and supplementary motor areas (SMA), insula and posterior parietal cortex as sensitive to expected effort costs (Croxson et al. 2009; Burke, Brünger, et al. 2013; Meyniel et al. 2013; Skvortsova et al. 2014). Together, effort-based valuation studies have consistently emphasized the relationship of ACC and VS to effortful behavior, but inconsistently present several possible loci for the integration of effort and value.

Effort relates to the difficulty of performing a behavior, such as physical force or response rate required within a limited time. Previous reports typically presented effort costs within passive or forced choice paradigms that demanded invigoration to meet upcoming effort challenges (Croxson et al. 2009; Prévost et al. 2010; Kurniawan et al. 2013). This is a distinct process from the prospective imposed cost of effort upon the subjective value of a potential outcome. Accordingly, recent reports suggest that ACC and VS activations in effort-based tasks may reflect these invigoration demands, rather than the valuation processes (McGinty et al. 2013; Holec et al. 2014; Shenhav et al. 2014). Similarly, single neurons within ACC were shown to exhibit encoding of task demands more often than effort-discounted value (Hosokawa et al. 2013). To specifically target effort- and risk-cost valuation, we developed a paradigm that presented participants with choices about effort and risk separated from effort production and outcome resolution. Participants were first trained to associate cues with physical grip force effort levels. In a subsequent choice phase, participants rated the subjective attractiveness of mixed gambles that offered a monetary gain or loss from their initial endowment. The outcome of these mixed gambles could depend upon future effort performance. Participants separately made similar choices in a task with fixed risk prospects. We hypothesized that prospective effort would impose a cost upon decision makers, reflected in both their willingness to gamble and neural activation related to value. Neuroimaging analyses examined the relationship of neural activity during choice to the magnitude of potential gain, loss and effort or risk present in each gamble. We further separately tested neural correlates of behavioral model-based predictions of subjective value from each task. Thus, we present a novel approach to characterize the neural basis of valuation under prospective effort, a common cost in economic choices.

We share all neuroimaging and behavioral data as well as analyses codes used.

## Materials and Methods

### Participants

Forty-six healthy, right-handed participants recruited from the University of Texas at Austin community participated in the experiment. Six participants were excluded from further analyses due to either excessive head movement, MRI artifacts or failure to meet behavioral criteria described below. The remaining forty participants comprised the sample group for our fMRI analyses (mean age = 22.62 years, standard deviation = 2.90 years, 21 females). Sample size was determined a priori by a power analysis for a contrast of interest (parametric prospective effort) from a pilot study of 13 participants (mean age = 21.3 years, standard deviation 2.31 years, 7 females) with the fMRIpower software package (Mumford and Nichols 2008). Each participant provided informed consent prior to the experiment. All participants had normal or corrected-to-normal vision, reported no history of psychiatric diagnoses, and neuralgic or metabolic illnesses. The Institutional Review Board at the University of Texas at Austin approved all experimental procedures.

### Behavioral paradigm

#### General methods

Prior to the experiment, participants were endowed with $20 cash. Participants were instructed that they were participating in a study about preferences in economic choices. Stimuli presentation and participant response collection were implemented with custom MATLAB code and the Psychophysics toolbox (Brainard 1997).

#### Baseline force measurement

Once inside the MRI scanner, each participant’s maximum voluntary contraction (MVC) force was assessed with an MR-compatible dynamometer (BIOPAC TSD121B-MRI, BIOPAC Systems Inc., USA). Participants were prompted to squeeze the dynamometer with their right hand as hard as possible in three intervals of two seconds interspersed with periods of rest for two seconds (Fig.1A). The calibration procedure was performed without feedback or incentives. The average force assessed by this procedure was considered the participants’ MVC for calibration of the training and test task phases. After calibration, participants were shown a display with real-time force feedback for demonstration purposes.

**Figure 1.**
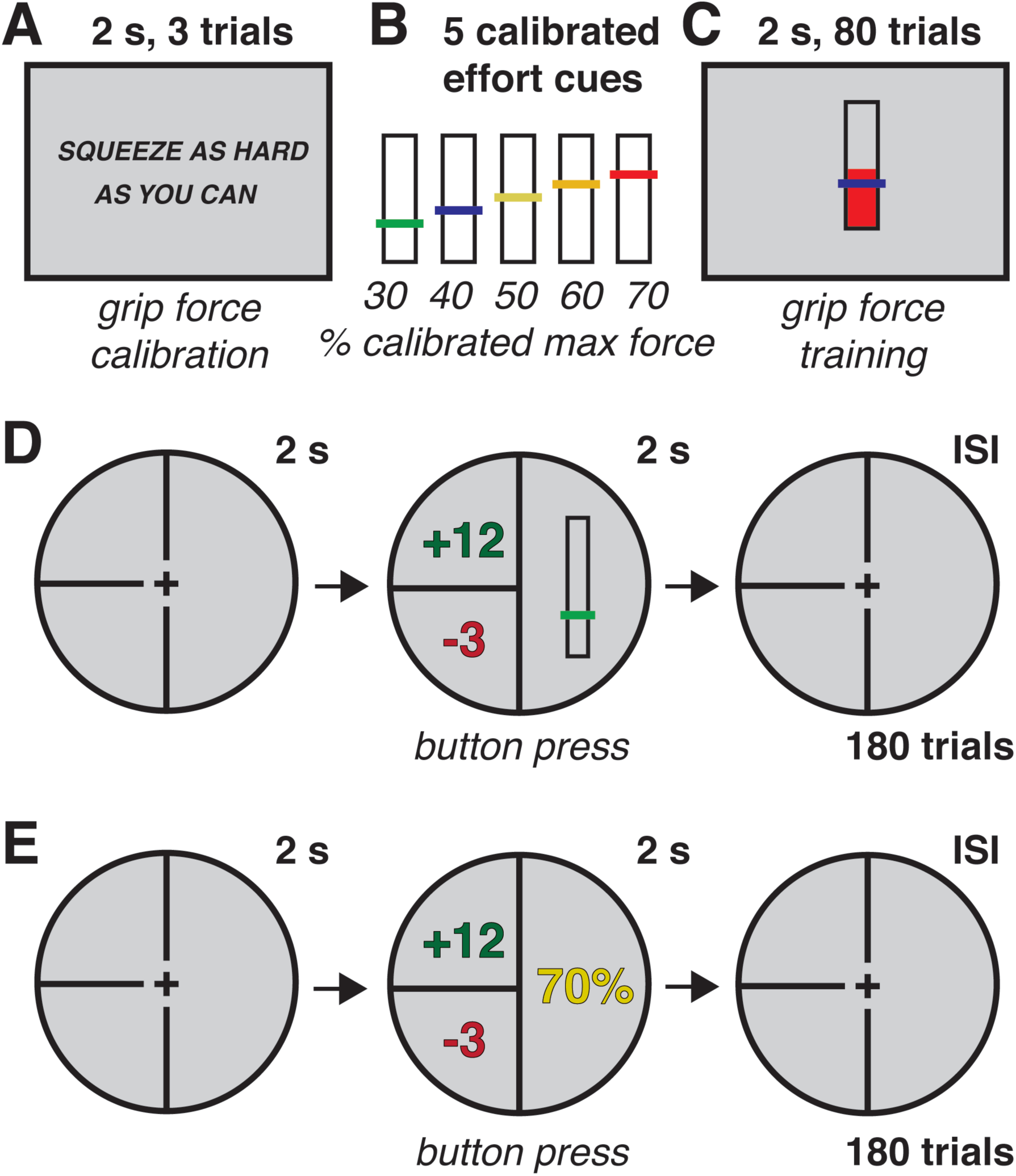
Summary of behavioral task: **A**, Effort level calibrated for each subject without incentive or feedback **B**, Training phase tested performance at 5 calibrated levels of grip force with real-time effort feedback. After training phase, subjects rated subjective attractiveness of prospective effort mixed gambles **C**, then prospective risk mixed gambles **D**, before resolution of one randomly selected gamble from each session.

#### Effort training

To examine the role of effort in economic choices, participants were first trained to associate color and height cues with the performance of physical grip effort levels. For each participant, five effort levels were determined at 30, 40, 50, 60 and 70% of their calibrated MVC (Fig.1B). During the effort training phase, participants attempted to complete physical grip effort production trials without incentives at five levels of difficulty. Each training run consisted of 80 total trials, 16 trials at each effort level, drawn randomly from one of five orders optimized for event-related fMRI. To successfully complete a trial, participants were required to exert grip force at or above a given effort level for at least 1 second of total time within a 2 second response period. Participants received two types of feedback during the training phase: real-time effort production level (dynamically adjusted height of red bar in effort display, Fig. 1C) and success or failure to meet the effort goal on a trial-by-trial basis.

#### Prospective effort mixed gambles

Following the effort training session, participants rated the attractiveness of prospective effort mixed gambles. Each mixed gamble presented participants with three components: a potential gain in addition to their endowment ($2-$12 in $2 increments); a potential loss from their endowment ($1-$6 in $1 increments) and one of the five effort levels established in the previous training session (30, 40, 50, 60 and 70% of calibrated MVC force; Fig. 1D). There were 180 trials in the choice set, all 36 combinations of potential gain and loss at each of the five effort levels. Effort level trial order and inter-stimulus interval timing were determined by an efficiency calculation for the prospective effort contrast of interest (Kao et al., 2009). Potential gain and loss amounts were drawn from the choice set for each effort level randomly without replacement. To encourage participants to reflect on the subjective attractiveness of mixed prospects rather than a fixed decision rule (e.g. accept all gambles with potential gain > $6), we asked participants to indicate one of four responses to each gamble (strongly accept, weakly accept, weakly reject, and strongly reject) as quickly as possible with a four-button response box. Participants were instructed that one gamble trial would be drawn at random at the end of the experiment. If the gamble was accepted, the resolution of the gamble (monetary gain or loss) would depend upon successful performance of the indicated effort level five times in succession without error at the end of the experiment as practiced during the earlier training session. If the randomly selected gamble trial was previously rejected, then no gain or loss would occur to their endowment.

#### Prospective risk mixed gambles

To compare effort-based valuation with valuation under risk, participants completed a similar set of prospective risk mixed gambles. This phase always followed the effort mixed gambles phase to limit potential framing effects (e.g. thinking of prospective efforts in terms of fixed probabilities). Prospective risk gambles also presented participants with three components: a potential gain in addition to their endowment ($2-$12 in $2 increments), and a potential loss from their endowment ($1-$6 in $1 increments), and one of five winning probability levels (10, 30, 50, 70 and 90%; Fig. 1E). The five levels were selected to span a wide range of success probabilities. Trial orders and choice sets were constructed similarly to the previous phase for the prospective risk contrast of interest. Participants indicated one of four responses as above to each trial and were informed that one trial from this phase would be selected at random at the end of the experiment and its outcome honored according to their choice to accept or reject, as in the previous phase.

#### Confidence ratings

To assess the effect of the grip force effort training paradigm on subsequent prospective effort gamble valuations, participants were asked to rate how confident they were that they could perform five successive grip force trials, as in the training phase, at nine levels (10–90% of their MVC in 10% increments). Participants used a button pad to freely indicate between 0 and 100% confidence in their ability to successfully perform a grip force at each effort level. This phase was always completed after both effort and risk mixed gamble phases to limit probability-based framing effects during prospective gambles valuations.

#### Resolution

Finally, while still in the scanner, participants faced the resolution of each prospective gamble phase. One trial from the prospective effort mixed gambles phase and one trial from the prospective risk mixed gambles phase, were randomly selected. For each selected trial, if the participant indicated ‘strong reject’ or ‘weak reject’ to the randomly drawn gamble, the associated gamble resolution phase ended without gain or loss. However, if for the selected trial the participant responded with ‘strong accept’ or ‘weak accept’, the drawn gamble was played out as follows: For the selected trial from the prospective effort mixed gambles phase, participants were required to attempt five consecutive physical grip force trials at the effort level of the randomly drawn prospect (e.g. 40% in Fig.1C). For the selected trial from the prospective-risk mixed gambles phase, participants passively viewed the outcome of the randomly drawn gamble played out by the computer. For each case, success resulted in the monetary gain associated with the drawn gamble and failure resulted with the associated loss. All monetary gains or losses that incurred in the resolution phase were added or subtracted from the participant’s initial endowment. After the experiment, participants were debriefed and paid in cash for their time and the results of the mixed gamble resolutions.

#### Behavioral analyses

We tested participants’ overall willingness to gamble (proportion of gambles accepted) prior to other behavioral analyses. This test revealed two participants that failed to meet a predetermined behavioral threshold (accepted or rejected more than 95% of all gamble trials) and excluded them from further analyses. From the training phase, we calculated mean success rate (percentage of making the goal in effort trials). We fitted a binary-logistic mixed-effects regression model to each of the mixed gambles phases to predict gamble acceptance by prospective effort or risk, gain and loss magnitudes. In addition, we fitted a linear model to predict acceptance rates across the different gamble components and reaction times. Finally, mean confidence rates were calculated from the confidence rating phase. All statistical analyses were performed in R using lme4 package.

### Neuroimaging

#### Image acquisition

Imaging data were acquired on a 3 Tesla Skyra MRI scanner system (Siemens) with a 32-channel head coil. Anatomical images for registration to Montreal Neurological Institute (MNI) template space were acquired with a high-resolution magnetization prepared rapid gradient echo (MPRAGE) pulse sequence (TR = 1900 ms, TI = 900 ms, TE = 2.43 ms, flip angle = 9°, FOV = 256, voxel size 1.0 × 1.0 × 1.0 mm). Functional images were acquired with a T2% weighted multiband echo-planar imaging sequence (TR= 1000 ms, multiband acceleration factor= 4, iPAT parallel acceleration factor = 2, TE=30 ms, flip angle = 63°, FOV=230, voxel size 2.4 × 2.4 × 2.4 mm; Moeller et al., 2010). For functional scans, fifty-six gapless axial slices were positioned 30° off the anterior commissure-posterior commissure line to reduce frontal signal dropout (Deichmann et al. 2003). Higher-order shimming was used to reduce susceptibility artifacts.

#### fMRI data preprocessing

Raw DICOM data images were converted to NIFTI format and organized to conform to the ‘Brain Imaging Data Structure’ specifications (BIDS; Gorgolewski et al., 2016). Preprocessing was conducted using FMRIPREP (version 1.0.0-rc13; Esteban et al., 2018; K. Gorgolewski et al., 2011, 2017). Within the FMRIPREP framework, each T1 weighted volume was corrected for bias field using N4BiasFieldCorrection v2.1.0 (Tustison et al. 2010) and skullstripped using antsBrainExtraction.sh v2.1.0 (using OASIS template). Cortical surface was estimated using FreeSurfer v6.0.0 (Dale et al. 1999). The skullstripped T1w volume was coregistered to skullstripped ICBM 152 Nonlinear Asymmetrical template version 2009c (Fonov et al. 2009) using nonlinear transformation implemented in ANTs v2.1.0 (Avants et al. 2008). Functional data was motion corrected using MCFLIRT v5.0.9 (Jenkinson et al. 2002). This was followed by co-registration to the corresponding T1-weighted volume using boundary based registration 9 degrees of freedom – implemented in FreeSurfer v6.0.0 (Greve and Fischl 2009). Motion correcting transformations, T1 weighted transformation and MNI template warp were applied in a single step using antsApplyTransformations v2.1.0 with Lanczos interpolation. Three tissue classes were extracted from T1w images using FSL FAST v5.0.9 (Zhang et al. 2001). Voxels from cerebrospinal fluid and white matter were used to create a mask in turn used to extract physiological noise regressors using a CompCor (Behzadi et al. 2007). Mask was eroded and limited to subcortical regions to limit overlap with gray matter, six principal components were estimated. Frame-wise displacement (Power et al. 2014) was calculated for each functional run using Nipype implementation. Spatial smoothing of functional images was performed with a Gaussian kernel with a full-width half maximum of 5 mm. Data and design matrices were high-pass filtered with a Gaussian-weighted least-squares straight line fit with a cutoff period of 100 s. Grand-mean intensity normalization of each functional image volume’s entire four-dimensional data set was performed by a single multiplicative factor.

### fMRI data analyses

#### Whole-brain analyses

All neuroimaging analyses focused on neural activity at the time of prospective valuation. In each whole-brain analysis, general linear models assessed the relationship of neural activity to mixed gamble variables or subjective value model predictions. The first whole-brain parametric model (GLM1) modeled five parametric regressors, three from the prospective effort mixed gambles phase: monetary gain, monetary loss and physical-demand (grip force effort level). To account for the uncertainty component of the effort phase (probability of not making the goal), two additional regressors modeled actual success-probability (goal success rate during training), and perceived success-probability (post task confidence rating). A second whole-brain parametric model (GLM2) modeled three parametric regressors from the prospective risk mixed gambles phase: monetary gain, monetary loss and prospective risk level associated with gamble resolution. In both models, parametric mixed gamble regressors were chosen to identify brain regions wherein activation or deactivation correlated with magnitude of each regressor, independent of changes in the other regressors without orthogonalization (Mumford et al. 2015). A second set of analyses (GLM3) and (GLM4) modeled a parametric regressor that reflected an estimate of a participant’s probability of accepting a gamble from the effort- and risk-based phases, respectively. For each participant, responses were collapsed into accept or reject categories and a binary logistic regression was fitted to predict response by the size of the potential gain, loss and effort (GLM3) or risk (GLM4). We then applied the resulting logistic equation parameters to generate a predicted probability of gamble acceptance regressor for each trial for each phase. In a separate whole brain-analyses, these regressors were used to identify brain regions that tracked the predicted probability of accepting a gamble, an estimate related to the subjective value of the gamble at choice. The binary logistic regression equations followed the forms:

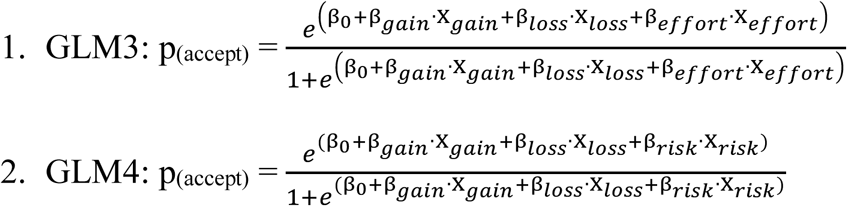

In all of the GLMs participants’ reaction times were modeled using the mean centered trial-by-trial reaction time. Additional regressors of no interest included all trials unmodulated (intercept), trials without behavioral responses, and nine confound regressors derived from preprocessing with FMRIPREP. These nine regressors included six motion parameters over time (rigid-body x-y-z translation and rotation transform) and three derivative of RMS variance over voxels (standardized, non-standardized and voxel-wise standardized). All statistical analyses were corrected for multiple comparisons with Gaussian Field Theory (GFT) based whole-brain cluster correction at a threshold of p <0.05.

### Data sharing

Unthresholded whole-brain statistical maps of activations are available for reference at NeuroVault.org (https://neurovault.org/collections/3955/; K. J. Gorgolewski et al., 2015). Neuroimaging data necessary to recreate all analyses will be made available as part of the OpenfMRI project (https://openneuro.org/datasets/ds001167/versions/00001; Poldrack & Gorgolewski, 2017). Behavioral data, analysis codes and fMRI analysis codes as well are available at: osf.io/mb3qw.

## Results

### Behavioral Results

#### Behavior: effort training

Participants’ behavior reflected modulation by effort, whereby success rates on effort training trials and post task confidence ratings decreased as the level of effort increased (Fig. 2A). In the training phase, increased effort requirements reduced the probability of successful performance (OR = 0.92, 95% CI [0.91, 0.93], p < 0.001). Averaging training performance within each effort level for each participant showed a strong negative linear correlation of effort level and training success ratio: (r = −0.81, t_(159)_ = −10.12, p < 0.001). Similarly, post-task confidence ratings of effort performance decreased as effort increased (r = −0.50, t_(319)_ = -17.95, p < 0.001); Fig. 2A).

**Figure 2.**
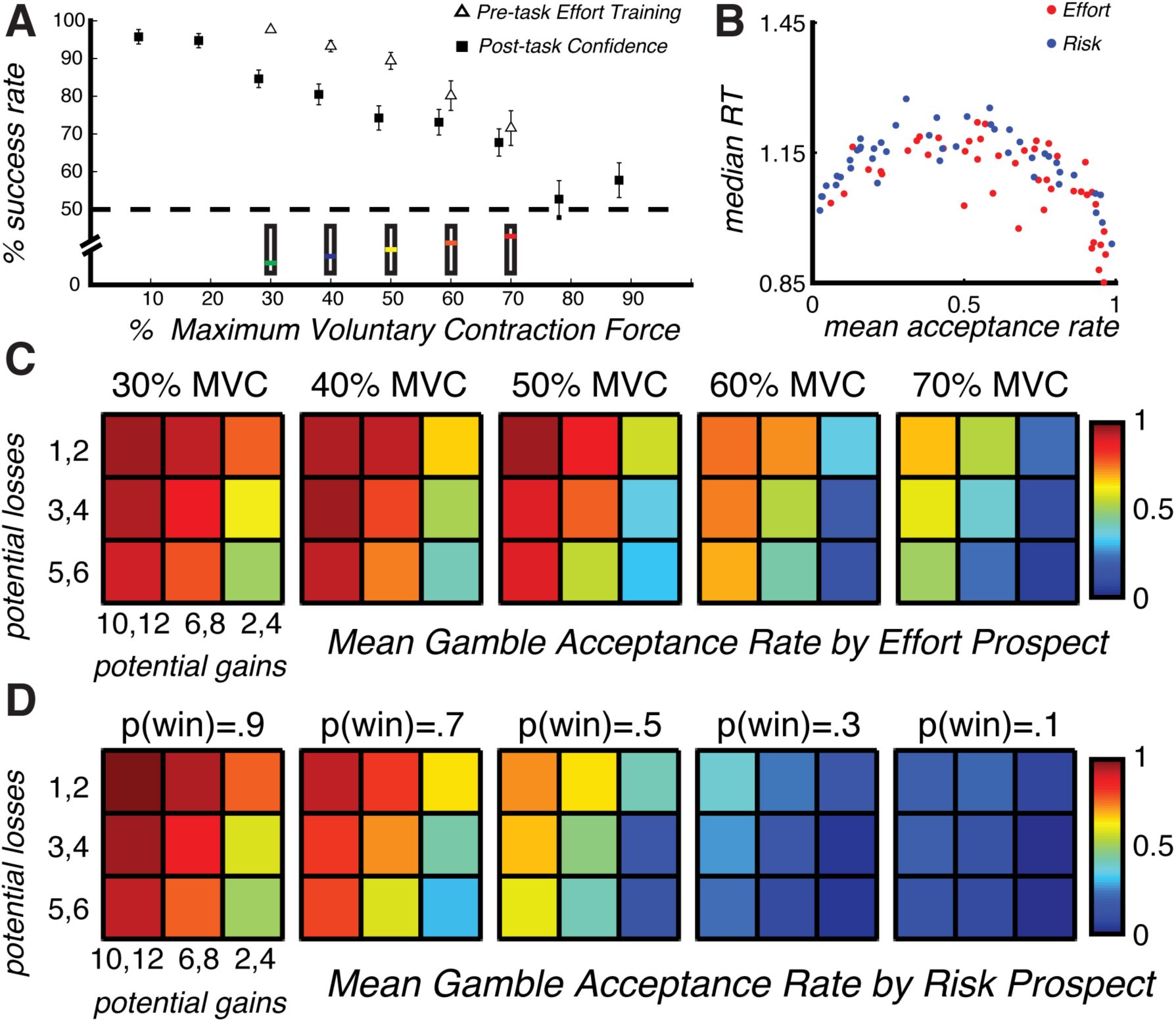
behavioral results: A. Increasing effort levels reduced performance success rate during training and confidence success rate during post-task survey. B. Group normalized median reaction times related to distance from indifference reflected in mean gamble acceptance rates in effort (red) and risk (blue) tasks. C,D. Heat maps depict preferences by level of prospective cost. For illustration, gain and loss were collapsed into 3 levels each. Error bars indicate S.E.M.

#### Behavior: mixed gambles

Participants’ overall willingness to gamble reflected an effect of prospective effort and risk costs (see Table 1). In the mixed gambles effort task, as the level of prospective effort increased, probability of gamble acceptance decreased. Similarly, in the mixed gambles risk task, prospective risk reduced average gamble acceptance rates.

**Table 1.** Prospective mixed gambles results

|  |  | **OR** | **95% CI** | **p** |
| --- | --- | --- | --- | --- |
| Prospective effort mixed gambles | Effort | 0.32 | [0.30, 0.35] | < 0.001 |
|  | Gain | 3.62 | [3.35, 3.90] | < 0.001 |
|  | Loss | 0.56 | [0.52, 0.60 ] | < 0.001 |
| Prospective risk mixed gambles | Risk | 0.11 | [0.10, 0.12] | < 0.001 |
|  | Gain | 3.27 | [3.00, 3.57] | < 0.001 |
|  | Loss | 0.51 | [0.48, 0.55 ] | < 0.001 |

Participants’ median reaction times followed a negative quadratic relationship (i.e. slowest median RT for gambles accepted on average 50% of the time) under prospective effort (R2 = 0.60, p < 0.001) and prospective risk (R2 = 0.78, p < 0.001; Fig. 2B). To illustrate participants’ choice preferences for mixed gambles under prospective effort and risk, gain and loss levels were collapsed into 3 bins each. Group average acceptance rates for effort-based gambles (Fig. 2C) and risk-based gambles (Fig. 2D) illustrate a pattern of decreased willingness to gamble across the entire choice set as prospective effort or risk increased.

### Neuroimaging results

#### Whole-brain parametric mixed gambles

Parametric analyses of neural activation at choice (GLM1, GLM2) revealed activity in distinct brain regions that was modulated by the components of effort- and risk-based mixed gambles tasks (Fig. 3). Increased activation for increasing prospective effort was found in left primary sensory cortex and increased activity for decreasing physical-demand was found in vmPFC (Fig. 3A). In contrast, increased activation for increased prospective risk showed a broader region that included left sensorimotor cortex and left superior parietal lobule. Decreased activation for increased prospective risk was found within the medial frontal cortex, anterior and posterior cingulate cortex, right striatum, frontal polar cortex and right sensorimotor cortex (Fig. 3D). In both the effort and risk tasks increasing prospective gain corresponded with increased activity in the right sensorimotor cortex, SMA, dorsal and posterior ACC and visual areas. In the effort task, increased activation for increasing prospective gains was also found within the ventral striatum, right hippocampus, ACC and ventral prefrontal regions (OFC, vmPFC Fig. 3B,E). In both effort and risk tasks, there were no decreased activations for increasing prospective gain. Under prospective effort, increased activations for increasing prospective loss were found in right dlPFC and lateral frontal polar cortex, left primary motor cortex, left posterior parietal cortex and medial premotor areas, and there were no decreased activations for increasing prospective loss (Fig. 3C). No activation were found for prospective loss under risk.

**Figure 3.**
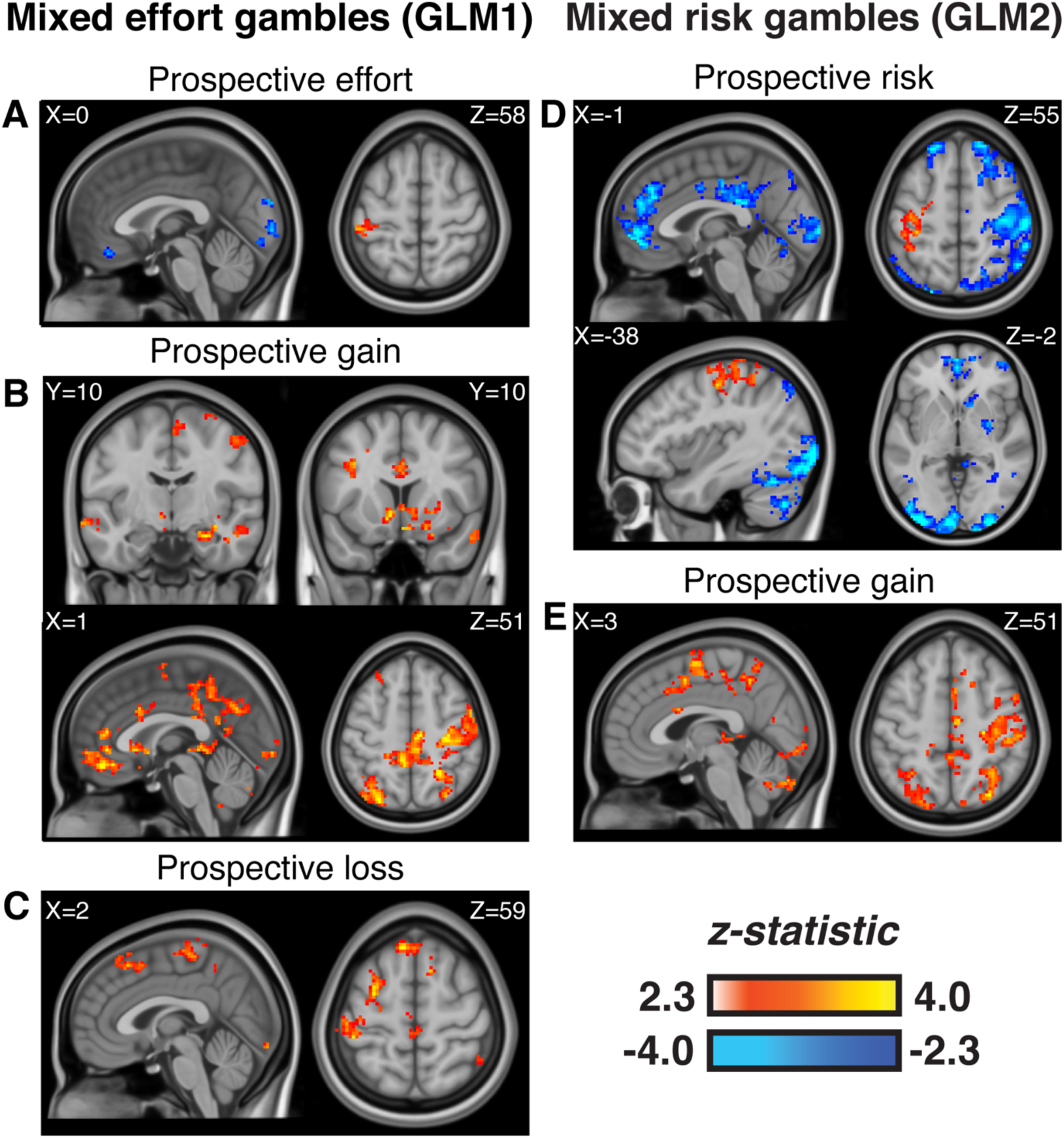
Parametric activation and deactivation related to magnitude of prospective effort **A**, potential gain **B**, and loss **C**, in effort-based mixed gamble stimuli, and activation related to prospective risk **D**, and gain **E**, in risk-based mixed gamble stimuli. Statistical maps corrected for multiple comparisons using Gaussian Random Field Theory at whole-brain level *p* < 0.05.

#### Whole-brain subjective value prediction analyses

A parallel set of parametric analyses (GLM3, GLM4) identified brain regions where activation tracked model predictions of the probability of accepting each gamble. In contrast to the parametric analyses that considered the fixed magnitude of mixed gamble components, these analyses relied upon each participant’s choice behavior to generate individual estimates of subjective value for each trial. These analyses revealed shared and distinct contributions of regions for valuation under each prospective cost. The right VS, vmPFC and right sensorimotor cortex tracked value estimates under both prospective costs. Under prospective effort, activity within left dlPFC, right insula and right putamen tracked value estimates (Fig. 4A). Under prospective risk, right lPFC, and bilateral caudate tracked value estimates (Fig. 4B).

**Figure 4.**
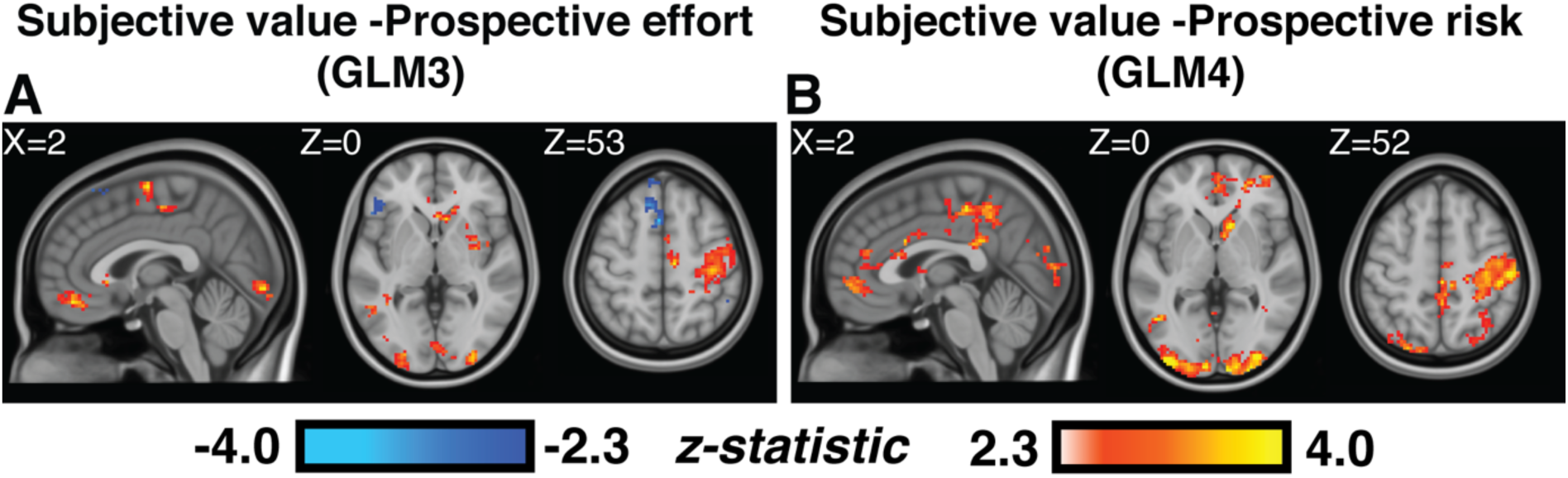
Parametric activation and deactivation related to estimates of subjective value from the effort **A**, and risk **B** tasks. Statistical maps corrected for multiple comparisons at whole-brain gaussian random field theory level *p* < 0.05.

## Discussion

We examined the neural basis of effort-based valuation isolated from production demands and outcome resolution. We adapted a prospective mixed gambles choice paradigm to present participants with mixed gambles associated with familiar effort challenges or fixed risk prospects. Participants’ choice behavior and confidence ratings separately reflected the imposition of a graded cost upon rewards associated with effort or risk, an observation consistent with many previous effort-based studies (Walton et al. 2006; Salamone et al. 2018).

Our neuroimaging analyses focused on the time of choice to reveal activation related to prospective effort or risk cost, and participants’ choice behavior. We found a modulation of vmPFC and left sensory cortex at choice by prospective effort magnitude. It has been suggested that vmPFC functions as a global comparator of multimodal valuation (Gläscher et al. 2009; Lebreton et al. 2009; Levy and Glimcher 2012), and several studies reported links between vmPFC activation and effort-based preferences (Basten et al. 2010; Treadway et al. 2012; Kroemer et al. 2014). Activity in primary sensory cortex has been reported to reflect the magnitude of effort demand (Keisker et al. 2009; Schmidt et al. 2012) as well as degree implied physical effort in read descriptions (Moody and Gennari 2010). Economic theories of decision making models often describe valuation as a cognitive process where options’ properties are compared independent of the sensorimotor contingencies (e.g., Padoa-Schioppa, 2011). In contrast, embodied cognition frameworks maintain that motor and sensory inputs are integral components of value representations (Kiefer and Pulvermüller 2012). Our findings support this interpretation of effort-cost representation as early as the relevant effector sensory cortex (hand area in our task), which in turn putatively relays the information for higher level integration in the vmPFC.

The present findings do not align with related literature that emphasizes the role of the ACC and VS in effort-based valuation processes (Prévost et al. n.d.; Rudebeck et al. 2006; Skvortsova et al. 2014; Klein-Flugge et al. 2016). Our results support recent neurophysiological findings that suggested that the contribution of ACC relates to anticipation or invigoration to effort demands, that are absent in a prospective task (Cowen et al. 2012; Holec et al. 2014). Critical reviews and reports put into question the link of dopamine-related activity within VS to effort and reward expectations (Salamone and Correa 2012; McGinty et al. 2013), and reframe the role of ACC in valuation as control-based (Shenhav et al. 2013). Likewise, related evidence suggests that ACC activation during foraging decisions reflect inferences of choice difficulty, not value (Shenhav et al. 2014). Furthermore, this interpretation is supported by similar neuroimaging studies that did not find significant evidence for ACC modulation by effort cost in prospective effort choice (Bernacer et al. 2016) and voluntary effort-based choices, either physical (Kurniawan et al. 2013) or cognitive (Schouppe et al. 2014),.

In addition to identifying regions that tracked effort cost, the goal of our analyses was to relate neural activation at choice to participants’ behavior. We used subjective responses to individually predict the probability of accepting each prospective mixed gamble, an estimate of its’ subjective value. We found that activation of vmPFC and VS as well as visual and motor regions tracked subjective value estimates during choices under both prospective effort and risk. Notably, clusters encompassing insula, putamen and dorsal premotor regions were related to subjective value estimates under prospective effort, whereas prospective risk-based estimates related predominantly to activation of bilateral caudate and right lPFC. These findings accord with evidence that vmPFC and VS comprise a core network for the computation of value during choice. In contrast to ACC, premotor areas and insular cortex that encode arousal or salience (Bartra et al. 2013; Rangel and Clithero 2013). Similar to other studies of effort-based decisions (Burke, Brunger, et al. 2013; Mathar et al. 2016), our design included an effort task where the participants did not reach the target goal in all of the effort trials. We accounted for this uncertainty component (probability of not making the goal) in our model, yet we acknowledge that future designs should separate between these components.

In summary, our results demonstrate the validity of the prospective effort paradigm and refine the association of several brain regions to the process of effort-based valuation that is independent of production. Notably, we provide evidence to support the notion that a common network represents value discounted by effort cost at choice as in other costs such as risk and delay (Kable and Glimcher 2007; Tom et al. 2007; Peters and Buchel 2009). While extensive literature supports a crucial role for ACC in effort-based valuation, our results suggest its contribution may be constrained to invigoration or anticipation of upcoming effort production. Given the great societal cost of maladaptive sensitivity to effort and its role in common impairments to motivated behavior, further understanding of the neural basis effort-based valuation offers great promise to inform the next generation of behavioral and neurological treatments.

